# Bacteriostatic antibiotics promote the evolution of CRISPR-Cas immunity

**DOI:** 10.1101/2021.05.04.442568

**Authors:** Tatiana Dimitriu, Elena Kurilovich, Urszula Lapinska, Konstantin Severinov, Stefano Pagliara, Mark D. Szczelkun, Edze R. Westra

## Abstract

Phage therapy can be used in combination with antibiotics to combat infections with bacterial pathogens^1–3^. However, bacteria can rapidly evolve phage resistance via receptor mutation, or using their CRISPR-Cas adaptive immune systems^4^, which insert short phage-derived sequences into CRISPR loci in the bacterial genome^5^ to guide sequence-specific cleavage of cognate sequences^6^. Unlike CRISPR-Cas immunity, mutation of the phage receptor leads to attenuated virulence when the opportunistic pathogen *Pseudomonas aeruginosa* is infected with its phage DMS3vir^7^, which underscores the need to predict how phage resistance evolves under clinically relevant conditions. Here, using eight antibiotics with various modes of action, we show that bacteriostatic antibiotics (which inhibit cell growth without killing) specifically promote evolution of CRISPR-Cas immunity in *P. aeruginosa* by slowing down phage development and providing more time for cells to acquire phage-derived sequences and mount an immune response. Our data show that some antimicrobial treatments can contribute to the evolution of phage-resistant pathogens with high virulence.

## Main text

We studied the effect of antibiotics on the evolution of CRISPR-Cas immunity in *Pseudomonas aeruginosa*, an opportunistic pathogen that commonly evolves antibiotic resistance and that has elicited large interest in phage and phage-antibiotic combination therapies for treatment of *P. aeruginosa* infections^8–10^. *Pseudomonas* strain PA14 can acquire immunity against its lytic phage DMS3vir^11^ by acquiring phage sequences (spacers) into CRISPR loci in its genome^12^. However, despite the presence of an active CRISPR-Cas system, this bacterium mostly evolves resistance through surface modification (SM) by loss or mutation of the type IV pilus (DMS3vir receptor) in rich broth and in artificial sputum medium that mimics the cystic fibrosis lung environment it commonly colonizes^7,13^.

### Antibiotics effects on phage resistance

To understand how antibiotics shape the population and evolutionary dynamics of *P. aeruginosa* during phage infection, we infected PA14 cultures grown in rich medium supplemented with sub-inhibitory concentrations of 8 different antibiotics (Extended Data Fig. 1A) with phage DMS3*vir*. Of these antibiotics, four are bactericidal (ciprofloxacin (Cipro), streptomycin (Strep), gentamycin (Gen) and carbenicillin (Carb)) and four are bacteriostatic (chloramphenicol (Chl), tetracycline (Tc) erythromycin (Erm), and trimethoprim (Tm)) against *P. aeruginosa* (Extended Data Fig. 1B). Most antibiotics delayed the phage epidemics and subsequently the evolution of phage resistance (Extended Data Fig. 2A and B). Nonetheless, at 3 days post infection (d.p.i.) phage resistance was essentially fixed in all cultures (Extended Data Fig. 2C). Strikingly, at this point, the type of phage resistance that had evolved was strongly dependent on the presence and the type of antibiotic. In the absence of antibiotics, or in the presence of bactericidal antibiotics, only a minority of bacteria evolved CRISPR-Cas immunity^13^, whereas a large proportion of the bacterial population evolved CRISPR-Cas immunity in the presence of bacteriostatic antibiotics (Fig. 1 and Extended Data Fig. 2D). This effect was relatively weak for Tm compared to Tc, Erm and Chl (Fig. 1). Chl was found to trigger CRISPR immunity across a wide range of concentrations, whereas no effect was observed when we used a Chl-resistant strain (Extended Data Fig. 3A). These data, and the fact that Chl, Tc, Erm and Tm have different modes of action, suggests that bacteriostatic antibiotics promote evolution of CRISPR immunity because they limit bacterial growth rates.

**Figure 1:**
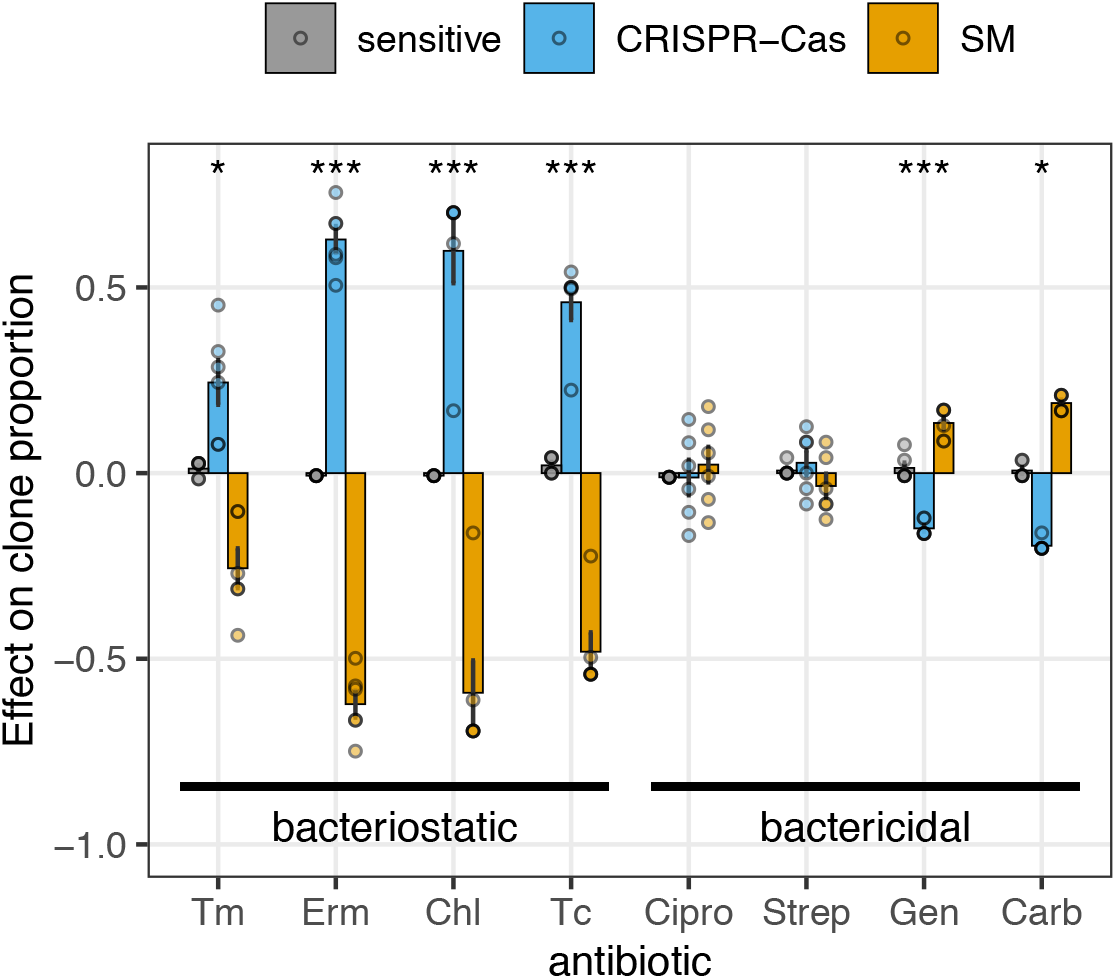
Bacteriostatic antibiotics promote CRISPR-Cas immunity. Effect of antibiotics on the proportion of sensitive (grey), CRISPR-Cas (blue) and SM clones (yellow) at 3 d.p.i., compared to the associated no-antibiotic treatment. Bars and error bars show mean ± s.e.m., and individual biological replicates are plotted as dots (N=6). Asterisks indicate antibiotics with CRISPR-Cas proportion significantly different from the associated no-antibiotic treatment (* 0.01<*p*<0.05; *** *p*<0.001). Antibiotics are ordered from left to right by decreasing minimum bactericidal concentration / minimum inhibitory concentration ratio, a measure of their bacteriostatic vs bactericidal activity. Raw data are shown in Extended Data Fig. 2D.

### Antibiotics effects on phage replication

The common feature of bacteriostatic antibiotics is that they inhibit cell growth, which might lead to higher rates of spacer acquisition^14^. To better understand the relationship between bacterial growth and evolution of CRISPR immunity, we first measured bacterial growth curves (based on the optical density, OD600, of the cultures) in the presence of each antibiotic at concentrations used in our evolution experiments. Analysis of exponential growth rates in batch culture and doubling times of individual cells in a microfluidics device^15–17^ showed that Chl and Tc cause particularly slow growth and this is associated with a large increase in the evolution of CRISPR-Cas immunity (Extended Data Fig. 4). More generally, this analysis revealed a correlation between exponential growth rate and the evolution of CRISPR immunity, with the exception of Erm, which was found to affect bacterial growth at higher cell densities that are closer to stationary phase.

Bacteriostatic and bactericidal antibiotics impact cell metabolism differently, and lead respectively to decreased and increased cell metabolic rates^18^. Because phage production is dependent on the metabolism and protein synthesis machinery of the host^19,20^, we hypothesized that bacteriostatic antibiotics may slow down phage replication, which could provide a larger window of time for the CRISPR-Cas immune system to acquire spacers from the phage prior to irreversible cell damage or cell death. To test this hypothesis, we performed one-step phage growth assays to detect when mature intracellular phages are produced. First, we pre-cultured cells for 12 hours in the presence of each antibiotic, so that cells were close to stationary phase and to the peak of phage epidemics in the absence of antibiotics. We found that all bacteriostatic antibiotics caused a strong delay in phage production compared to cells cultured in the absence of antibiotics (Fig. 2A). Under those conditions, we were unable to analyse the effect of bactericidal antibiotics on phage production, due to high rates of cell death in these treatments. Interestingly, when we next analysed the effects of antibiotics during exponential growth, we found that Erm had no effect on phage production (Fig 2B), consistent with its minor effect on exponential growth rate of the bacteria (Extended Data Fig. 4). All other bacteriostatic antibiotics (Chl, Tc and Tm) delayed the formation of infectious progeny phages (Fig. 2B), and this effect was observed across a broad range of Chl concentrations (Extended Data Fig. 3B). These data strongly suggest that inhibition of bacterial growth by bacteriostatic antibiotics causes a delay in the phage eclipse period. Bactericidal antibiotic had more variable effects: Cip and Carb showed no interference with production of infectious phages, in agreement with their mechanism of action and known synergy with phage therapy^21^, whereas the presence of Strep or Gen (both aminoglycosides) resulted in very little phage production (Fig. 2B) due to a large proportion of unproductive infections (Fig. 2C). Interestingly, we also observed a small but significant increase in failed infections in the presence of Erm and Tc. While this may contribute to the evolution of CRISPR immunity^22^, it is insufficient to explain the effects of bacteriostatic antibiotics in general, since these had only small (Erm and Tc) or no effects (Tm, Chl) on the proportion of unproductive infections (Fig. 2C).

**Figure 2:**
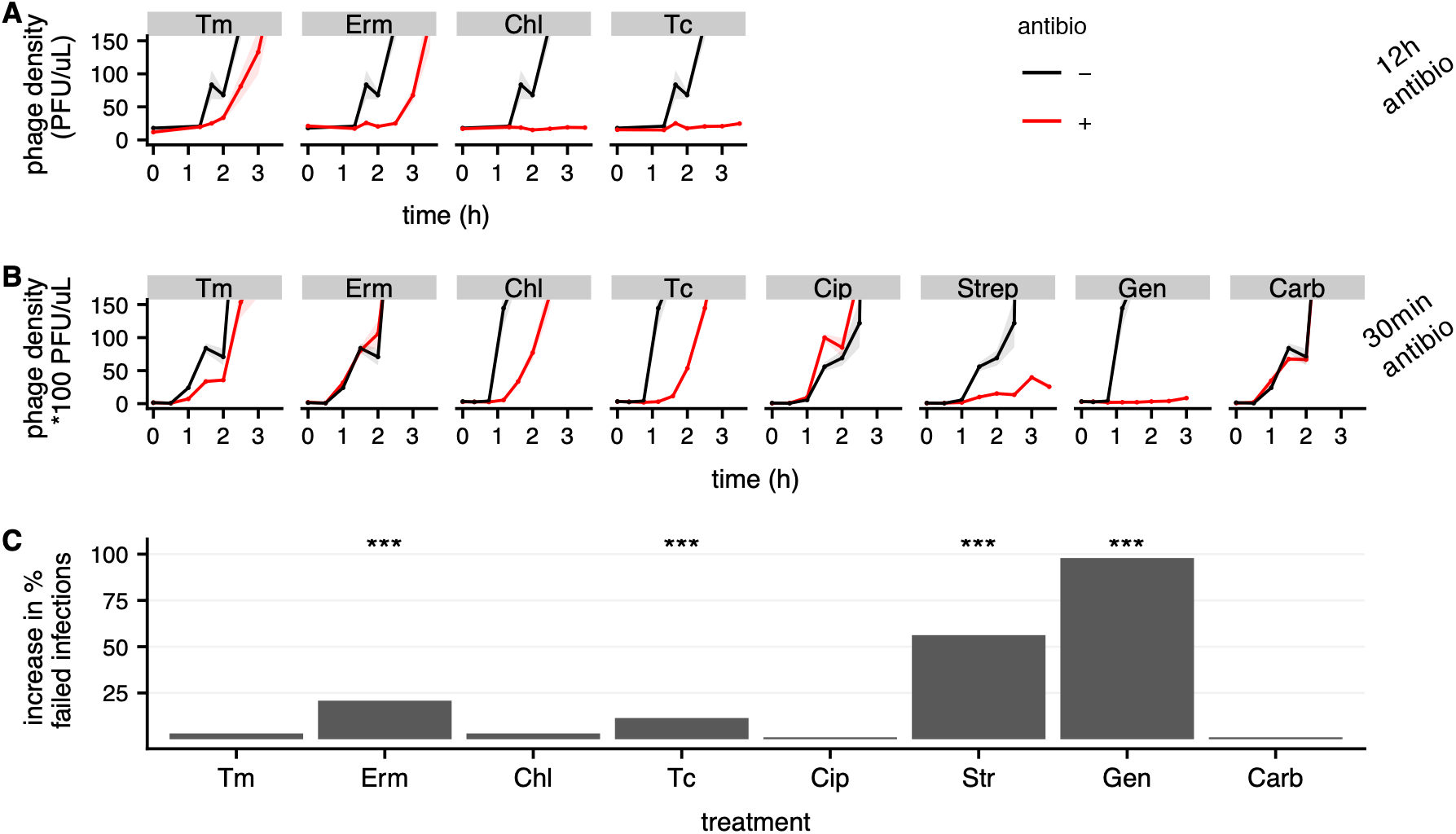
Bacteriostatic antibiotics delay production of mature phage particles. A and B: Effect of antibiotics on phage production dynamics. Phage density over time during infection of cells that are pre-exposed to antibiotics for 12h (A) or 30min (B) are shown in red and no-antibiotic controls in black. Lines and shaded areas are respectively mean and s.e.m. (N=4). C: Effect of antibiotics on the frequency of failed phage infections. Barplots show the increase in percentage of infected populations in which no phage was detected after 24h. Asterisks indicate antibiotics with a significant increase in the number of populations with no phages (chi-square tests, *** *p*<0.001).

### Antibiotics effects on spacer acquisition

We hypothesized that the reduced rate of within-host replication of phage in the presence of bacteriostatic antibiotics may provide more time for infected cells to acquire CRISPR-Cas immunity (i.e. to insert phage-derived spacers into CRISPR loci on the bacterial genome). To test this hypothesis, we performed short-term (3h) infection assays and measured the proportion of bacteria that had acquired new spacers in the presence or absence of each antibiotic. This revealed that in the presence of all bacteriostatic antibiotics more cells acquire CRISPR-Cas immunity, whereas bactericidal antibiotics had no effect (Fig. 3A). This effect was detectable despite bacteriostatic antibiotics inhibiting absolute cell growth (Extended Data Fig. 5). Across antibiotics, the rate at which CRISPR immunity is acquired in these short-term experiments was significantly correlated to the levels of CRISPR-Cas immunity that evolved at 3 days post infection (Fig. 3B, Pearson’s correlation t_1,6_=3.9, *p*=0.008, ρ=0.85). Crucially, none of the antibiotics except Cip affected the rates at which bacteria with SM are generated (Extended Data Fig. 6). Furthermore, none of the antibiotics caused an increase in Cas protein production (Extended Data Fig. 7). Finally, we also tested whether antibiotics impact the way selection acts on clones with CRISPR immunity and receptor mutants. For example, CRISPR immunity may be more effective (and therefore provide a greater fitness advantage) when phage replicates more slowly. However, competition experiments between a clone with CRISPR-Cas immunity (CR) and a SM-resistant clone showed that the presence of bacteriostatic antibiotics had either no impact or reduced the fitness of CRISPR immune bacteria relative to receptor mutants (Extended Data Fig. 8). Collectively, these data therefore show that bacteriostatic antibiotics increase the frequency at which cells survive phage infection as a result of the acquisition of CRISPR-Cas immunity when phage replicate more slowly. Consistent with this conclusion, the bacteriostatic antibiotic Chl only triggered increased evolution of CRISPR immunity if it was present during the first day following phage infection (Extended Data Fig. 9), when most cells are still phage-sensitive (Extended Data Fig. 2C), and later exposure, when bacteria have already acquired resistance, had no effect.

**Figure 3:**
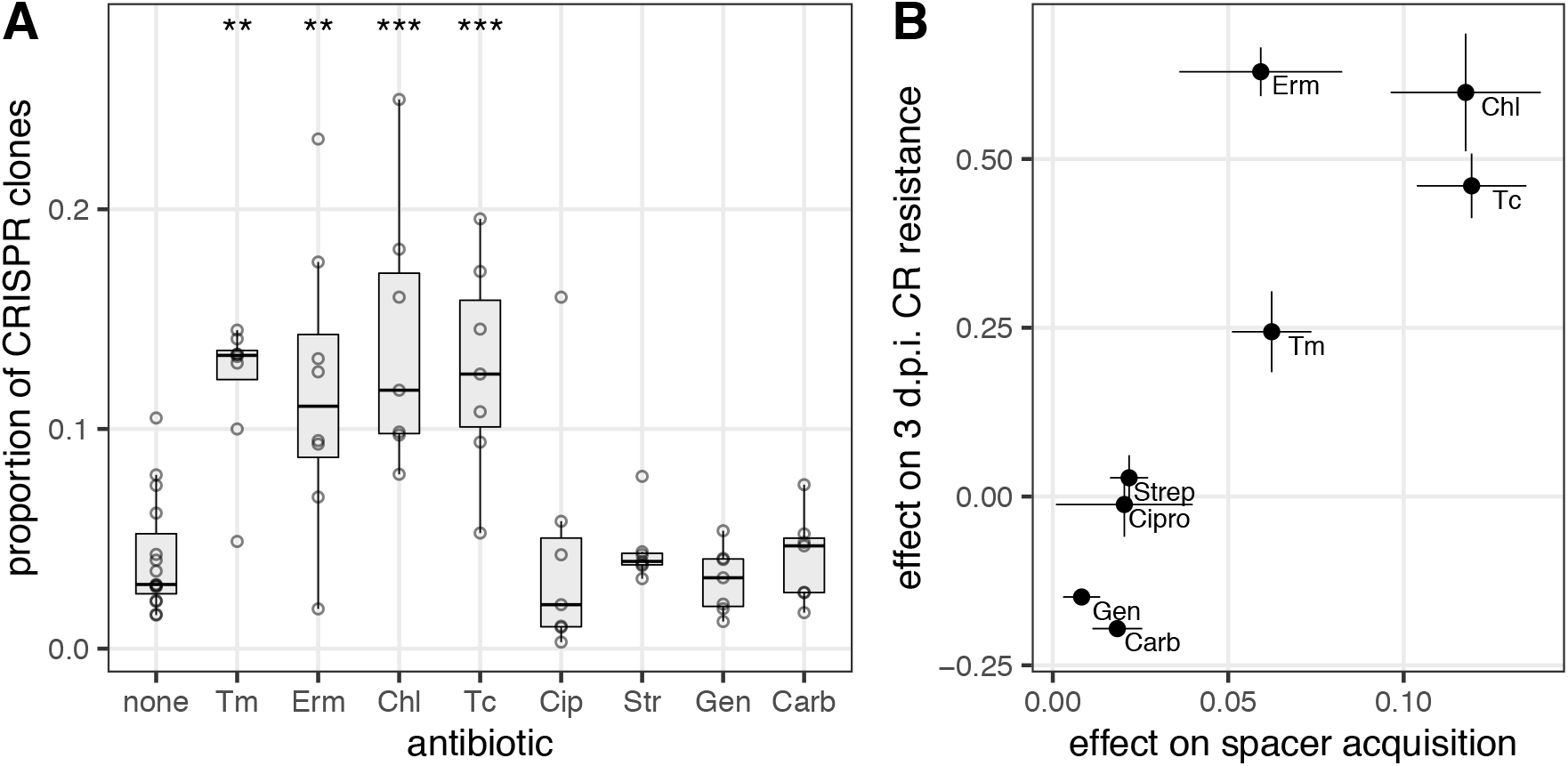
Bacteriostatic antibiotics increase acquisition of CRISPR immunity. A shows the proportion of resistant clones which are CRISPR-Cas immune after 3h phage infection. The centre value of the boxplots, boxes and whiskers respectively represent the median, first and third quartile, and 1.5 times the interquartile range; dots show individual data points (N=6). Asterisks show treatments significantly different from the no-antibiotic control (Tukey HSD; **, 0.001<*p*<0.01; *** *p*<0.001). In B, the average change in proportion per treatment is plotted against the average increase in proportion of CRISPR-Cas immune clones in the evolution experiments shown in Fig. 1, error bars showing s.e.m. (N=6).

### Poor nutrients also promote CRISPR-Cas

Given that bacteriostatic antibiotics promote the evolution of CRISPR immune bacteria, we hypothesized that other environmental factors that slow down bacterial growth rate could also lead to an increase in the evolution of CRISPR immunity. To test this hypothesis, we performed evolution experiments in minimal medium with different carbon sources (glycerol, glucose and pyruvate), which are associated with different maximum growth rates (Extended Data Fig. 10A). One-step growth curves in these different media showed that phage production was delayed when cells grew on the poorest carbon source, glycerol (Extended Data Fig. 10B), as expected based on changes in cell metabolic rates^20,23,24^. Infection experiments at two different phage inoculum sizes, which maximizes the dynamic range of any changes in the evolution of CRISPR-Cas immunity^13^, showed that carbon source had a significant effect on the proportion of bacteria that acquired CRISPR-Cas immunity (Extended Data Fig. 10C, F_2,32_=16.8, *p*=10^−5^). Specifically, cells growing on glycerol grew more slowly and acquired CRISPR-Cas immunity more frequently than those growing on glucose (Tukey HSD, *p*=0.038), whereas cells growing on pyruvate grew more rapidly and acquired CRISPR-Cas immunity less frequently (Tukey HSD, *p*=0.008). This was not due to differences in overall phage production^13^ (Extended Data Fig. 10D and E) and therefore suggests that bacterial growth rate is a key determinant of the frequency at which bacteria acquire CRISPR-Cas immunity.

## Discussion

Phage therapy can benefit patients by killing bacteria and by exploiting trade-offs between phage resistance and pathogen virulence^25^ or antibiotic resistance^2,26^. However, *P. aeruginosa* clones that acquire CRISPR-Cas immunity can escape these trade-offs and retain virulence^7^. Previous studies have identified a number of environmental variables that shape the evolution of CRISPR immunity by affecting the fitness of CRISPR-Cas immune *P. aeruginosa* clones relative to those with mutated phage receptors^7,13,27^. Here, we identify clinically relevant environmental factors which increase the frequency at which sensitive *P. aeruginosa* clones acquire CRISPR immunity during a phage infection. Acquisition of CRISPR-Cas immunity relies on the acquisition of spacers from infecting phages, subsequent expression of appropriate protective CRISPR RNAs and interference. It is a major limiting step because cells that just acquired spacers might still be killed before mounting a sufficient immune response^5,28^. We found that bacteriostatic antibiotics promote the acquisition of CRISPR-Cas immunity over a large range of concentrations, due to slowing bacterial growth, which in turn delays phage development. This is consistent with increased spacer acquisition from plasmids under conditions that are associated with slow bacterial growth^14,29^. Interestingly, a number of CRISPR-Cas systems have recently been found to induce dormancy following target recognition^30–32^ or to be coupled to genes that induce dormancy^33,34^. A dormancy response of infected cells with CRISPR immunity can benefit neighbouring cells by eliminating phage from the environment and by limiting the invasion of phage mutants that overcome CRISPR immunity^31^. Our data suggest the possibility that another advantage of a dormancy response could be that they may lead to more efficient spacer acquisition during infections. Our findings may also help to explain why the acquisition of CRISPR-Cas immunity is relatively rare under laboratory conditions, in which bacteria commonly grow at rates much higher than in the wild^35^, and in clinical contexts. *Pseudomonas aeruginosa* displays slow growth rates in biofilms^36^ and in cystic fibrosis^37^, which will be compounded by antimicrobial treatment. Our results suggest that phage-antibiotic combination therapy should consider the possibility of increased evolution of CRISPR immunity.

## Methods

No statistical methods were used to pre-determine sample size. The experiments were not randomized, and investigators were not blinded to allocation during experiments and outcome assessment.

### Bacteria, phages and growth conditions

Except when stated otherwise, evolution experiments and phage assays used *P. aeruginosa* UCBPP-PA14 (PA14) and lytic phage DMS3vir^11^. UCBPP-PA14 *csy3::lacZ* was used for phage stock amplification, phage titre determination and estimation of Cas protein expression. Competition experiments used a surface mutant (3A) derived from PA14 *csy3::lacZ* and a CRISPR-resistant mutant (BIM4, bacteriophage insensitive mutant with 2 additional acquired spacers against DMS3vir) derived from PA14, both of which have been previously described^38^. DMS3vir and a mutant expressing anti-CRISPR against PA14 IF system, DMS3vir *AcrIF1*, were used for determination of resistance phenotypes^13^. Evolution experiments in the presence of Chl also used PA14-*cat*, a Chl-resistant mutant of PA14 carrying the *cat* gene inserted into the genome using a variant of plasmid pBAM^39^ carrying *cat*. To this end, the *cat* gene was amplified from plasmid pKD3^40^ using primers TAGATTTAAATGATCGGCACGTAAGAGGTT and CTGACCCTTGTCTTACGCCCCGCCCTGCCACT, then ligated into pBAM1 after digestion with SwaI and PshAI. For microfluidics experiments, we used PA14 *flgK::Tn5B30(Tc*^*R*^)^41^. All bacterial strains were grown at 37 °C in LB broth or M9 medium (22 mM Na_2_HPO_4_; 22 mM KH_2_PO_4_; 8.6 mM NaCl; 20 mM NH_4_Cl; 1 mM MgSO_4_; and 0.1 mM CaCl_2_) supplemented with 40 mM glucose, glycerol or pyruvate. All liquid cultures were grown with 180 rpm shaking. For experiments using M9, overnight pre-cultures were themselves grown in M9 with the same carbon source.

### Determination of antibiotic activity

For MIC (minimum inhibitory concentration) determination, overnight cultures (∼5.10^9^ cells/mL) were diluted 10^4>^-fold in LB medium. 20 μL of the diluted cultures were inoculated into 96-well microplate wells containing 180 μL of LB supplemented with antibiotics using 2-fold serial dilutions of the antibiotic. After 18 h growth at 37 °C, MIC was determined as the lowest antibiotic concentration with no visible growth. To determine the MBC (minimal bactericidal concentration), the content of wells with no visible growth was plated on LB-agar and further incubated overnight. MBC was defined as the lowest antibiotic concentration resulting in 99.9% decrease in initial inoculum cell density (< 5 CFU in 100 μL). MBC/MIC ratio was used to estimate if antibiotic activity was bacteriostatic or bactericidal: a high MBC/MIC ratio indicates that the concentration sufficient to prevent growth is much lower than the concentration required to kill the majority of cells^42^. In our assay, antibiotics with average MBC/MIC ratio >1 were the ones that are commonly recognized as being bacteriostatic (Tm, Erm, Chl and Tc).

### Evolution experiments

Evolution experiments were performed in glass vials containing 6 mL growth medium and appropriate antibiotics at the concentrations shown in Extended Data Fig. 1. 60 µL from overnight cultures were co-inoculated with 10^4^ plaque-forming units (p.f.u.) of phage DMS3vir, with the exception of the experiment in Extended Data Fig. 10, where two different phage inocula of 10^4^ and 10^9^ p.f.u. were used. 1:100 volume was then transferred every 24 h into fresh medium for 3 days with the exception of the experiment in Extended Data Fig. 10, which was carried out for 5 days. Each treatment contained 6 biological replicates. Cell densities and phage titers were monitored daily with serial dilution in M9 salts (after chloroform treatment for phages), and enumeration of colonies on LB-agar and enumeration of plaques on a lawn of PA14 *csy3::lacZ* cells. The identification of phage resistance type (sensitive, CRISPR-Cas or SM) was performed by cross-streaking 24 randomly selected colonies on DMS3vir and DMS3vir-AcrF1 phages: SM clones are resistant to both phages and have a characteristic smooth colony morphology, whereas clones with CRISPR-Cas immunity are resistant to DMS3vir but sensitive to DMS3vir-AcrF1^13^.

### Determination of bacterial growth rate by optical density

Overnight cultures were diluted 100-fold into fresh growth media. Growth of 200 μL of culture was measured in a 96-well plate by measuring optical density at λ=600nm (OD600) for 14 to 24 h at 37 °C in a BioTek Synergy 2 Plate reader, with 5 s shaking before each measurement. All growth curves were performed in at least 8 replicates. Exponential growth rate was determined in R using the package growthrates^43^.

### Determination of bacterial doubling time by microfluidics

The mother machine device was fabricated and handled as previously reported^15,16^. Briefly, overnight cultures in LB were spun down via centrifugation for 5 minutes at 4000 rpm at room temperature (Eppendorf 5810 R). The supernatant was filtered twice (Medical Millex-GS Filter, 0.22 µm, Millipore Corp.) and used to re-suspend the bacteria to an OD600 of 75. 2 µl of the bacterial suspension was injected into the microfluidic mother machine device and incubated at 37 °C until there were 1-2 bacteria in the lateral side channels. Fluorinated ethylene propylene tubing (1/32” × 0.008”) was connected to the inlet and outlet holes and connected to a computerized pressure-based flow control system (MFCS-4C, Fluigent) controlled by MAESFLO software (Fluigent) and outlet reservoir respectively. Spent media was flushed through the device to wash excess bacteria out of the main channel at 300 µL/h for 8 minutes to completely exchange the fluid in the device and tubing. The chip was mounted on an inverted microscope (IX73 Olympus, Tokyo, Japan) and images were acquired in bright-field via a 60×, 1.2 N.A. objective (UPLSAPO60XW, Olympus) and a sCMOS camera (Zyla 4.2, Andor, Belfast, UK) with a 0.03s exposure. The microfluidic device was moved by two automated stages (M-545.USC and P-545.3C7, Physik Instrumente, Karlsruhe, Germany, for coarse and fine movements, respectively) to image multiple fields of view in a sequential manner. The imaging setup was controlled by LabView. After acquiring the first set of images, we flowed each of the investigated antibiotics dissolved in LB at the appropriate concentration at 300 µL/h for 8 minutes before lowering the flow rate to 100 µL/h for 3 hours. The entire assay was carried out at 37 °C in an environmental chamber surrounding the microscope. Bacterial doubling times were extracted from the acquired image sets as previously reported^17^. Briefly, we tracked each individual bacterium and its progeny throughout each experiment and doubling times were measured as the lapses of time between successive bacterial divisions that were assessed by eye through the images loaded in ImageJ and considered to have happened when two daughter cells became clearly distinguishable from their respective parental cell.

### One-step phage growth assays

Overnight cultures of PA14 were first diluted into 6 mL growth medium ± antibiotic treatment in glass vials (N=4). For experiments with 30 min of pre-exposure to antibiotics, cells were diluted 25-fold into fresh media with antibiotics and grown for 30 min before phage addition. For experiments with 12 h of pre-exposure to antibiotics, cells were diluted 100-fold into fresh media with antibiotics and grown for 12 h before phage addition. Bactericidal treatments were excluded from further analysis because they caused a 4-fold to 570-fold decrease in cell density after 12 h, making it impossible to determine the latent period of phage under those conditions. After growing in the presence of antibiotics, approximately 5.10^7^ p.f.u. of DMS3vir were added in each vial, and vials were vortexed and incubated at 37 °C for 15 min, allowing phage adsorption. Cultures were then diluted 1000-fold into 6 mL growth medium ± antibiotic treatment to limit further adsorption and re-infection, vortexed again and transferred to 24-well plates for parallel processing. Samples were taken immediately (t=0) and then approximately every 20 minutes. The first samples were diluted in M9 salts and plated on LB-agar to quantify cell densities; all samples were chloroform-treated and plated on PA14 *csy3::lacZ* lawns. Phage densities measured after chloroform treatment correspond to the sum of free phages and mature phage particles inside infected cells.

### Determination of antibiotic effects on infection success

Four overnight cultures of PA14 were diluted in parallel 100-fold into LB with or without antibiotics. After 2 h growth at 37 °C, DMS3vir phages were added to a final concentration of 1000 p.f.u./mL (equivalent to 5 p.f.u. in 200 μL) and the vials were vortexed. After 15 min at 37 °C, vials were vortexed again and 24* 200 μL of each individual culture were aliquoted into 24 wells of a 96-well plate. Plates were incubated at 37 °C for 22 h, then 20 μL of each well were spotted on a lawn of PA14 *csy3::lacZ* cells in two replicates. With an average phage inoculum of 5 phages, the distribution of phages across wells is expected to follow a Poisson distribution with 0.7% wells containing 0 phages and 1.3% wells containing more than 10 phages. The control treatment with no antibiotics was consistent with this, as 1 in 96 wells produced no lysis. Lysis indicated that the founding phages reproduced. The number of wells in which phages failed to reproduce was counted for each treatment, and significance was determined by chi-square tests between antibiotic and no-antibiotic treatment.

### Measurement of mutation towards SM

To evaluate the frequency of SM cells in the absence of phage selection, cells were grown in LB ± antibiotic treatment for 24 h. After 24 h, cultures were serially diluted in M9 salts, then dilutions were plated both on LB-agar to calculate total cell density, and on LB-agar containing a high concentration of DMS3vir, which was generated by covering the agar surface with a phage stock of 10^8^ p.f.u./μL. The density of SM mutants was calculated from counting the number of colonies growing on top of DMS3vir. Three independent experiments were run with 6 experimental replicates each.

### Spacer acquisition assay

20 μL of PA14 overnight culture were first diluted 1:50 into 1 mL LB with or without antibiotics in 24-well plates, in 8 replicates per treatment. After 30 min of growth at 37 °C, 2.10^9^ DMS3vir phages were added per well, and cultures were incubated at 37 °C for 3h. The density of phage-sensitive cells was measured by plating 100 μL on LB-agar after 10^4^-fold dilution in M9 salts. The density of phage-resistant cells was measured by directly plating 100 μL of cultures on LB-agar without dilution: the phage density on these plates was sufficient to prevent growth of sensitive colonies. The majority of colonies had a smooth morphology characteristic of SM clones. We confirmed that smooth colonies were resistant to both DMS3vir and DMS3vir+AcrIF1, whereas non-smooth colonies were resistant to DMS3vir but sensitive to DMS3vir+AcrIF1, and were therefore CRISPR immune. In each culture, the proportion of CRISPR-Cas immune clones within the total population of resistant clones (CRISPR-Cas and SM) was calculated.

### Competition assays

Competition experiments were performed in 6 mL LB supplemented in the presence or absence of antibiotics. They were initiated by inoculating 60 μL of a 1:1 mix of LB overnight cultures of CRISPR-Cas immune (BIM2) and surface mutant (3A) clones. For treatments including phages, 8.10^9^ p.f.u. DMS3vir were added per vial. Samples were serially diluted at 0 and 24 h and plated on LB agar supplemented with 50 μg/mL X-Gal, to determine the ratio of the surface mutant that carries the *LacZ* gene and therefore forms blue colonies, and the BIM2, which forms white colonies. The selection rate of the CRISPR-Cas clone was calculated as m_BIM2_-m_3A_, with m the Malthusian parameter defined as log(density(t_1_)/density(t_0_)) ^44^. We used selection rate rather than relative fitness because some treatments led to an absolute decline in the abundance of the CRISPR-Cas clone.

### Cas expression assay

We used PA14 *csy3*::*lacZ* as a reporter strain for Cas gene expression, using the β-Galactosidase fluorogenic substrate 4-Methylumbelliferyl β-D-galactoside^45^ (MUG). An overnight culture of PA14 *csy3*::*lacZ* was diluted 100-fold into 6 mL of LB with or without antibiotics. After 5 h of growth, OD600 was recorded and 100 μL aliquots were immediately frozen at -80 °C. Prior to the assay, the frozen 96-well plate was defrosted and 10 μL were transferred to a new plate and frozen again at -80 °C for 1 h. After transfer to 37 °C, 100 μL reagent solution (0.25 mg/mL MUG and 2 mg/mL lysozyme in phosphate-buffered saline) was added to each well. Fluorescence was measured for 30 min in a Thermo Scientific Varioskan flash plate reader at 37 °C, with excitation and emission wavelengths respectively 365 nm and 450 nm. The 15 min timepoint was used for analysis. Relative fluorescence was calculated as (fluorescence at 15 min – fluorescence at 0 min) / OD600.

### Statistics

All statistical analyses were done with R version 3.4.1 ^46^, and package cowplot^47^. Evolution experiments with antibiotics were not all performed simultaneously (Figure 1, Extended Data Fig. 1, Extended Data Fig. 3): for these, we used individual Student t-tests comparing each treatment to the associated no-antibiotic treatment. For experiments in M9 that used two different phage inocula (Extended Data Fig. 10), the effect of the two treatments was modelled as: proportion of CRISPR-Cas immune clones ∼ carbon source + phage inoculum.

## Acknowledgments

This work was funded by grants from the European Research Council under the European Union’s Horizon 2020 research and innovation programme (ERC-2017-ADG-788405 to M.D.S and ERC-STG-2016-714478 to E.R.W.) E.R.W. was further supported by a NERC Independent Research Fellowship (NE/M018350/1).

## Author contributions

Conceptualization of the study was done by T.D. and E.R.W. Experimental design was carried out by T.D. Bacterial evolution, competition and growth experiments as well as phage infection assays were done by T.D. with assistance from E.K., and MIC measurements were done by E.K. Microfluidics experiments were designed and carried out by U.L. and S.P. Formal analysis of results was done by T.D. K.S. and M.D.S. contributed to discussions and provided feedback throughout the project. T.D. wrote the original draft of the manuscript, with later edits and reviews by T.D., K.S., S.P., M.D.S. and E.R.W.

