## Supplementary Figures for "Bacteriostatic antibiotics promote the evolution of CRISPR-Cas immunity"

**A**

| Abbreviation | antibiotic | target | MIC (μg/mL) | Concentration used | Static/cidal |
|---|---|---|---|---|---|
| Chl | Chloramphenicol | 50S ribosome subunit | 30 | 25 | Bacteriostatic |
| Tc | Tetracycline | 30S ribosome subunit | 10 | 2.5 | Bacteriostatic |
| Erm | Erythromycin | 50S ribosome subunit | 100 | 5 | Bacteriostatic |
| Tm | Trimethoprim | Dihydrofolate reductase | 6.25 | 2.5 | Bacteriostatic |
| Gen | Gentamycin | 30S ribosome subunit | 1.25 | 0.625 | Bactericidal |
| Str | Streptomycin | 30S ribosome subunit | 25 | 12.5 | Bactericidal |
| Carb | Carbenicillin | Penicillin binding proteins | 25 | 2.5 | Bactericidal |
| Cip | Ciprofloxacin | DNA gyrase | 0.2 | 0.1 | Bactericidal |

**B**

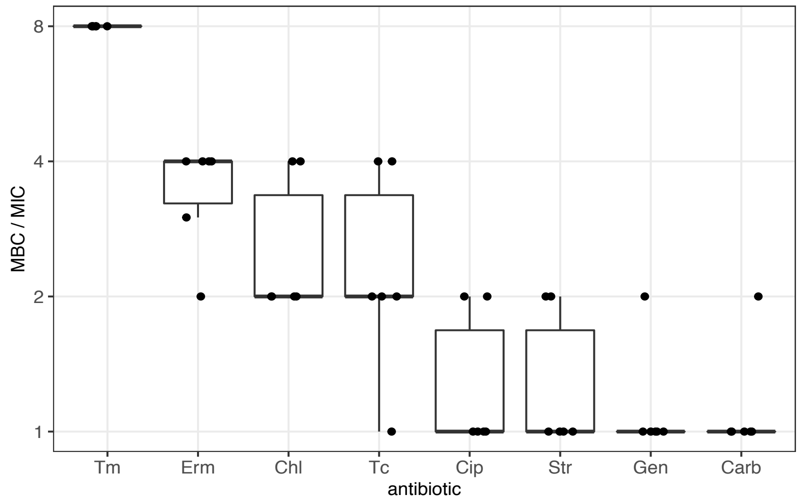


**Figure S1: Antibiotics used in this study and inhibitory effect.** A shows antibiotic molecular targets, standard concentrations used for experiments and overall bacteriostatic / bactericidal effect. MIC, minimum inhibitory concentration; MBC, minimum bactericidal concentration. In B, the MBC (minimum bactericidal concentration) vs MIC (minimum inhibitory concentration) concentration ratio is shown for all antibiotics tested. The centre value of the boxplots shows the median, boxes the first and third quartile, and whiskers represent 1.5 times the interquartile range; individual data points are shown as dots (N=8). Bacteriostatic antibiotics have on average MBC/MIC ≥2, whereas bactericidal antibiotics have MBC/MIC ~1.

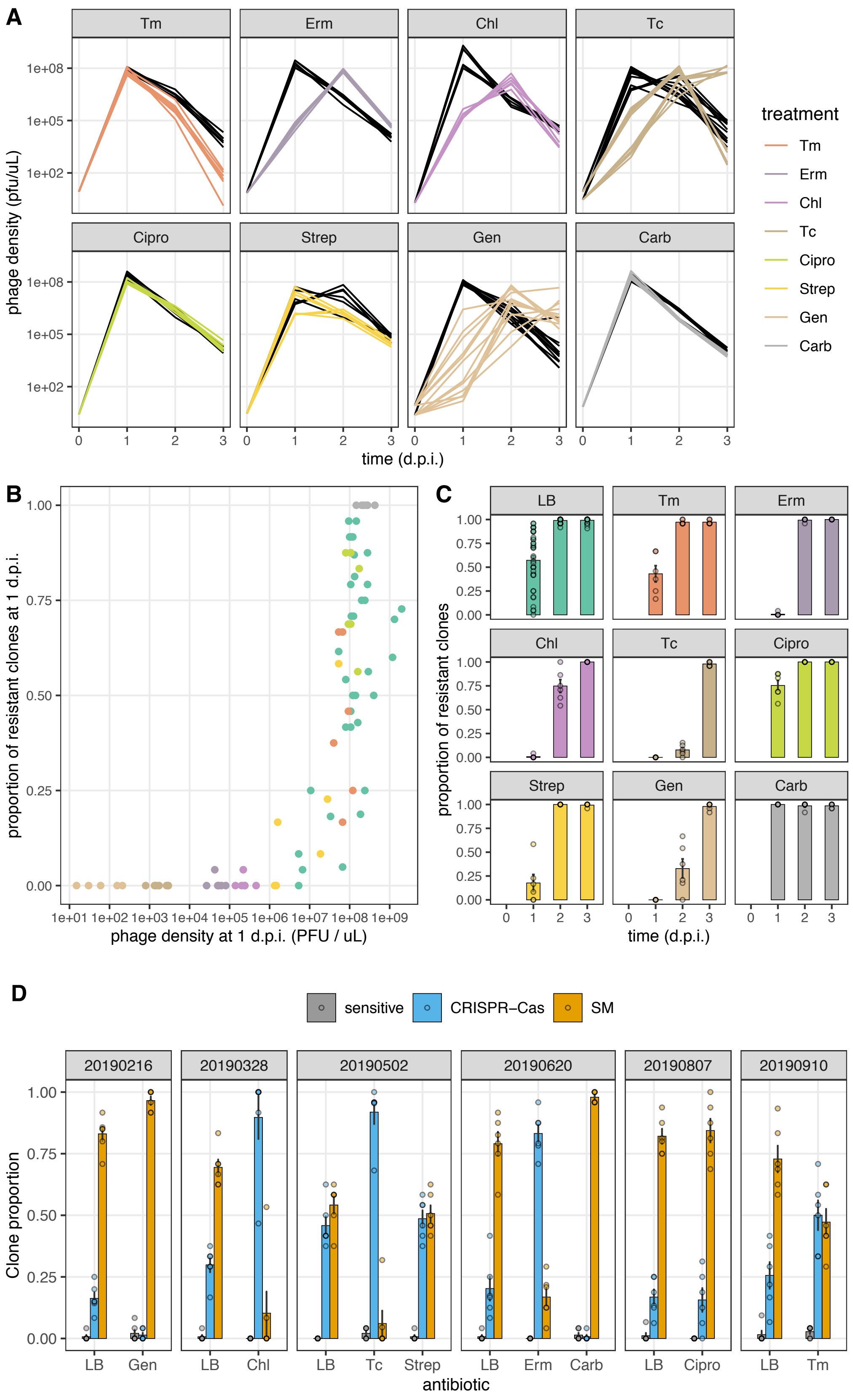
A
Tm
Erm
Chl
Tc
Cipro
Strep
Gen
Carb
phage density (pfu/uL)
1e+08
1e+05
1e+02
time (d.p.i.)
treatment
Tm
Erm
Chl
Tc
Cipro
Strep
Gen
Carb
B
proportion of resistant clones at 1 d.p.i.
phage density at 1 d.p.i. (PFU / uL)
1e+01 1e+02 1e+03 1e+04 1e+05 1e+06 1e+07 1e+08 1e+09
C
LB
Tm
Erm
Chl
Tc
Cipro
Strep
Gen
Carb
proportion of resistant clones
time (d.p.i.)
D
sensitive
CRISPR−Cas
SM
20190216
20190328
20190502
20190620
20190807
20190910
Clone proportion
0.00
0.25
0.50
0.75
1.00
LB
Gen
Chl
Tc
Strep
Erm
Carb
Cipro
Tm
antibiotic

**Figure S2: Phage and phage resistance dynamics during evolution experiments.** A: phage population dynamics. Densities of phages are shown as a function of time for each antibiotic treatment (in colour) and associated no-antibiotic control treatment (in black). B: proportion of clones resistant to DMS3*vir* at 1 d.p.i. as a function of the phage density measured at 1 d.p.i.. The proportion of resistant clones present after 1 day of infection is higher in populations in which phage spread rapidly during the first 24h. In A and B, individual replicates are shown (N=6 per treatment). C: proportion of clones resistant to DMS3*vir* per treatment over time. D: Detail of antibiotic effect on phage resistance phenotypes. Proportion of sensitive (grey bars), CRISPR-Cas (blue bars) and SM clones (yellow bars) at 3 d.p.i., in the absence (LB) or presence of antibiotic treatment, ordered by experiment. In C and D, bars and error bars show mean ± s.e.m., and individual biological replicates are plotted as dots (N=6).

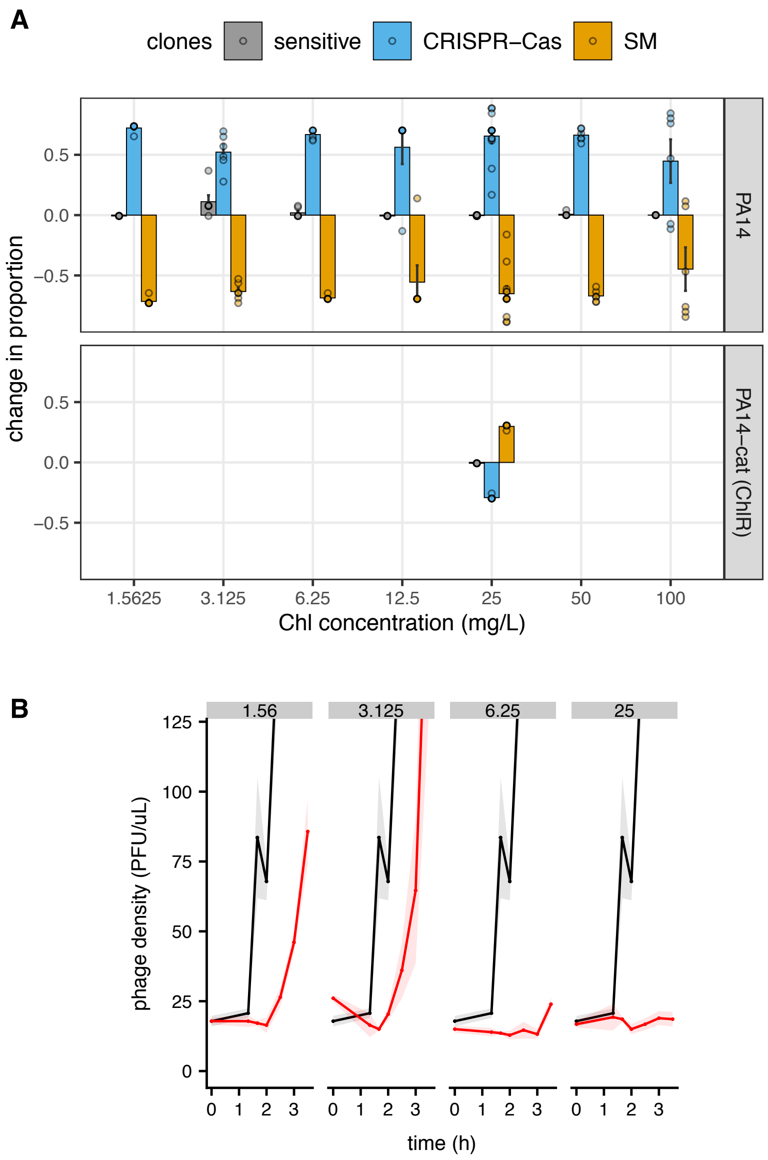


**Figure S3: Effect of a large range of Chl concentrations on CRISPR-Cas resistance evolution and phage replication dynamics.** A: Effect of Chl on the proportion of sensitive, CRISPR-Cas and SM clones at 3 d.p.i., compared to the associated no-antibiotic treatment. Bars and error bars show mean ± s.e.m., and individual biological replicates are plotted as dots (N=6). B: One-step phage growth assays in the presence of varying Chl doses. The no-antibiotic treatment is shown in black and antibiotic treatments in red, lines and shaded area are respectively the mean and s.e.m. (N=4).

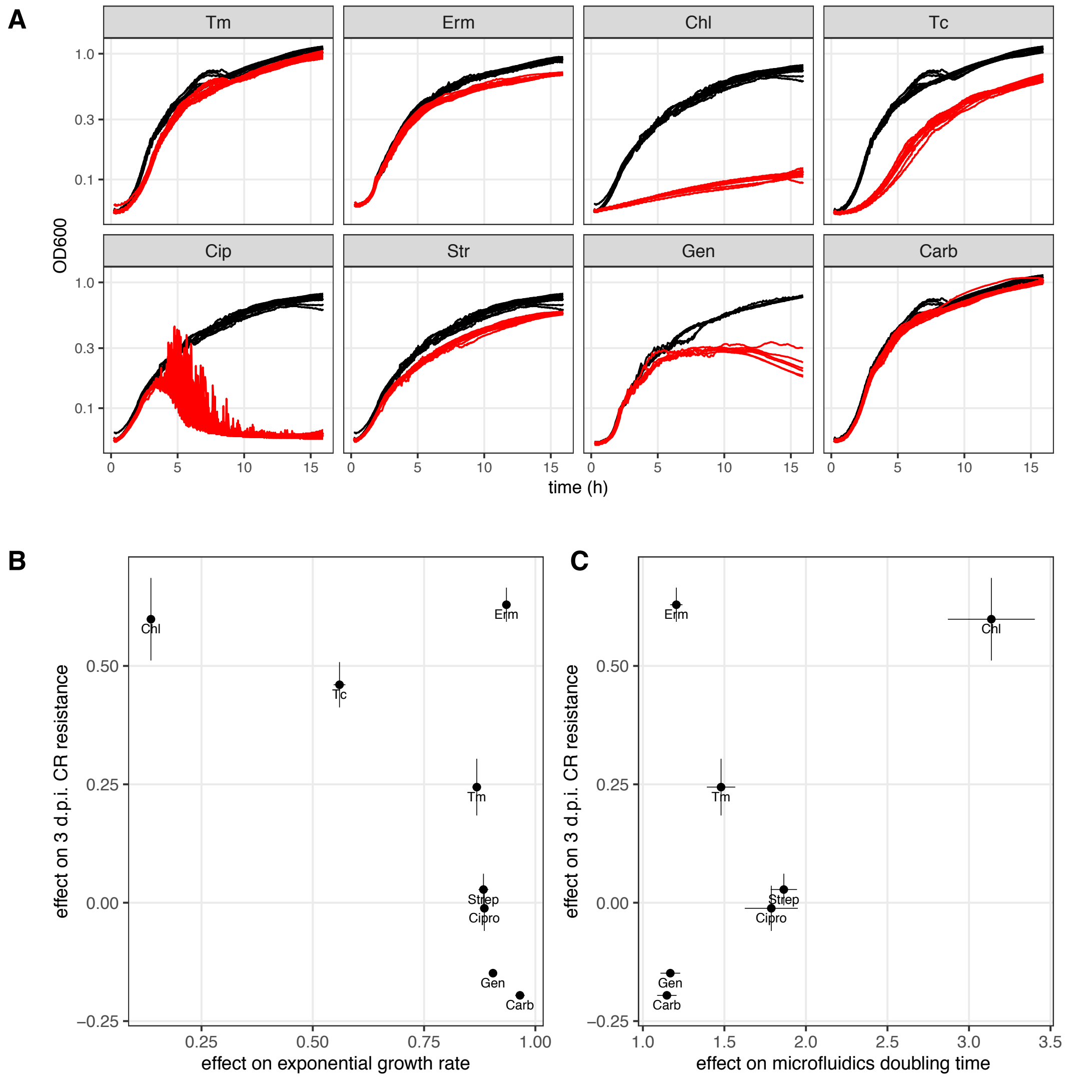


**Figure S4: Detail of antibiotic effect on cell growth.** In A, full OD600 growth curves are shown for all antibiotics (no antibiotic treatment in black, antibiotic treatment in red), at the antibiotic concentrations used for Figure 1. The aspect of the curves with Cip treatment was due to formation of cell aggregates. In B, antibiotic effect on 3 d.p.i. evolved CR resistance is shown as a function of exponential growth rate effect measured by OD600 change in 96-well plates (shown in A). In C, antibiotic effect on 3 d.p.i. evolved CR resistance is shown as a function of antibiotic effect on doubling time measured in microfluidics. Dots and error bars show respectively mean ± s.e.m. Antibiotic effect on 3 d.p.i. evolved CR resistance was not correlated to exponential growth rate (Pearson's correlation, $p$=0.11) or microfluidics doubling time ($p$=0.26) when taking all antibiotics into account, but was significantly correlated when excluding Erm, which acts closer to stationary phase (exponential growth rate, $p$=0.008, ρ=-0.89; microfluidics doubling time, $p$=0.018, ρ=0.89).

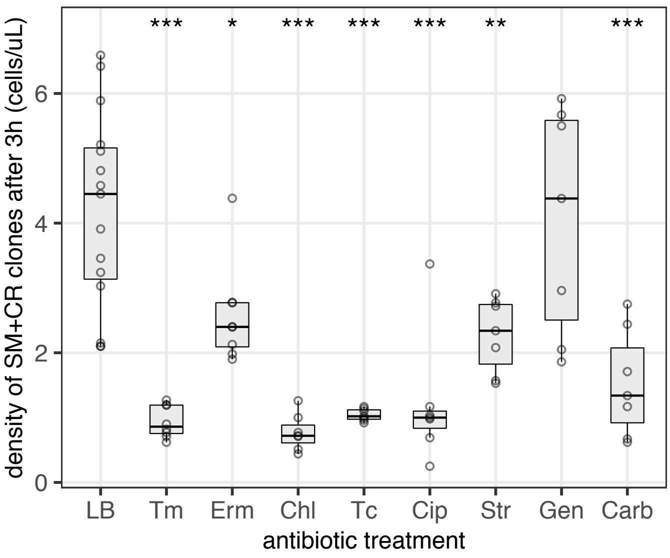


**Figure S5: Effect of antibiotics on short-term growth of PA14 phage-resistant clones.** The total phage-resistant cell density at the end of the spacer acquisition assay shown in Figure 3A. The centre value of the boxplots shows the median, boxes the first and third quartile, and whiskers represent 1.5 times the interquartile range; individual data points are shown as dots (N=6). Asterisks show treatments significantly different from the no-antibiotic control (*, 0.01<$p$<0.05; **, 0.001<$p$<0.01; *** $p$<0.001).

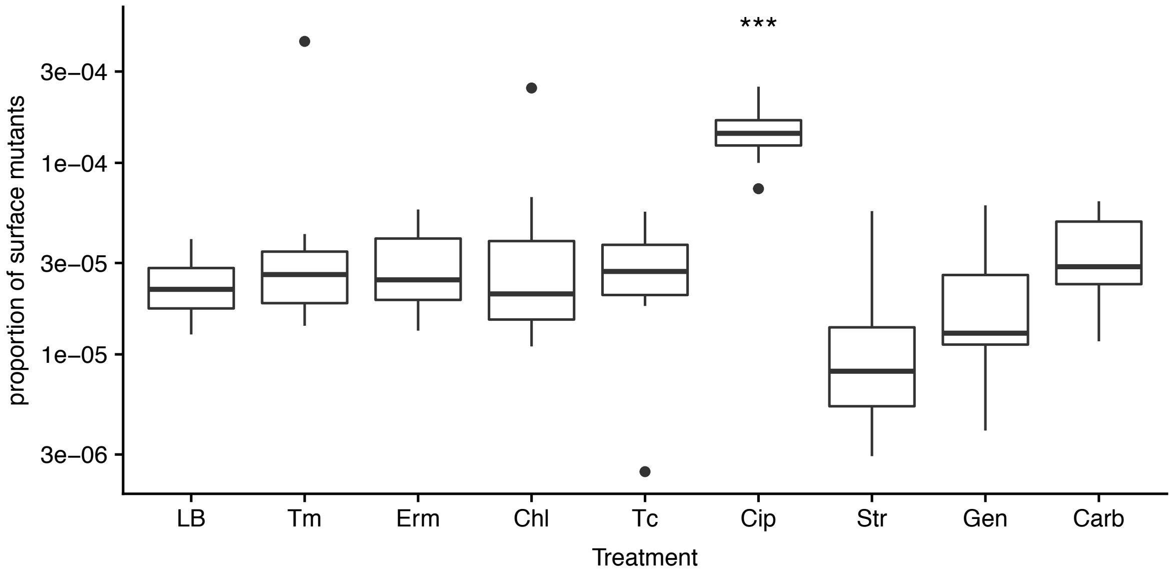


**Figure S6: Proportion of SM clones in populations grown in the absence of phage.** The centre value of the boxplots shows the median, boxes the first and third quartile, whiskers represent 1.5 times the interquartile range and dots are outliers (N=18 from 3 independent experiments). Asterisks show treatments significantly different from the no-antibiotic controls (***, $p<0.001$).

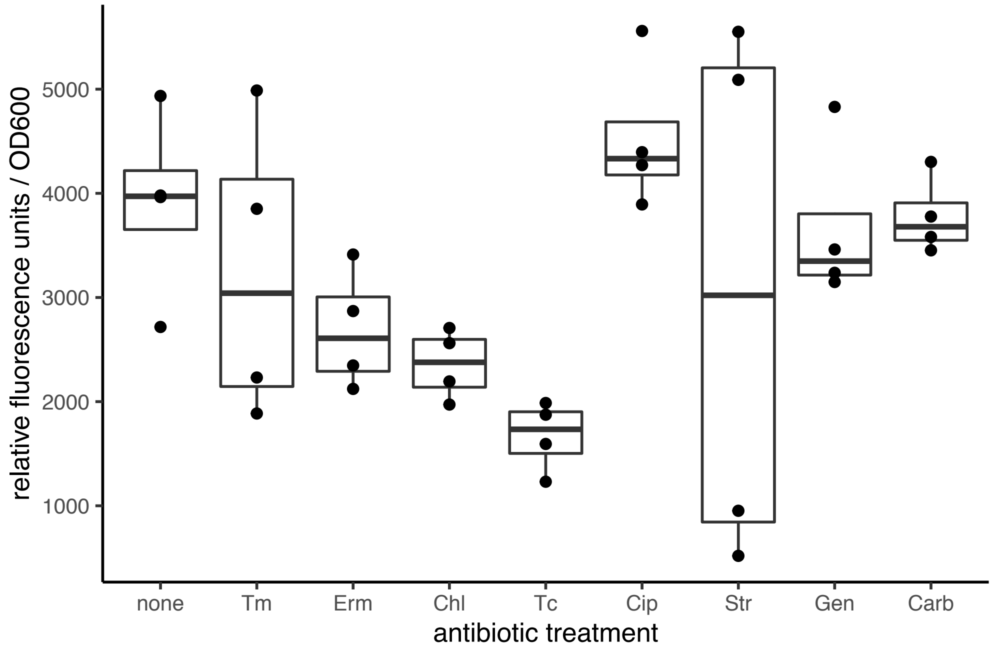


**Figure S7: Effect of antibiotics on Cas gene expression.** Relative fluorescence / OD600 is shown for the reporter strain PA14 csy3::lacZ grown for 5h with antibiotic treatment. The centre value of the boxplots shows the median, boxes the first and third quartile, and whiskers represent 1.5 times the interquartile range; individual data points are shown as dots (N=4). None of the antibiotic treatments had a significant effect compared to the no-antibiotic control (Tukey tests, all $p > 0.05$).

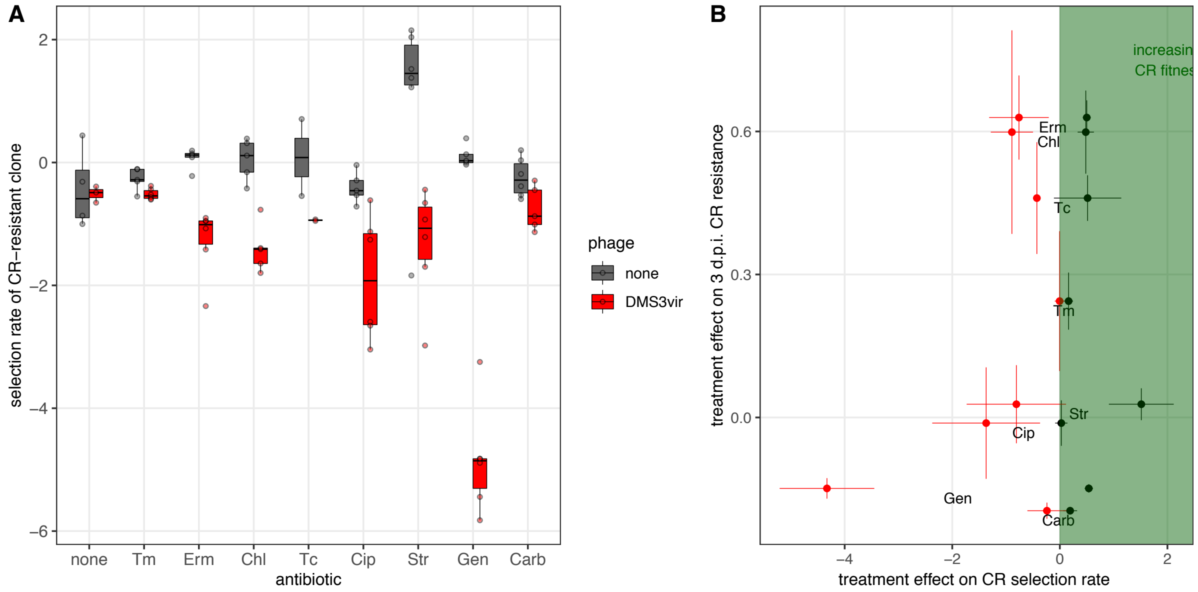

**Figure S8: Resistant clone fitness effects.** In A, the selection rate of the CRISPR-Cas resistant clone BIM2 against the surface mutant 3A is plotted as a function of antibiotic treatment, in the absence (gray) or presence (red) of phage DMS3vir). The centre value of the boxplots shows the median, boxes the first and third quartile, and whiskers represent 1.5 times the interquartile range; individual data points are shown as dots (N=6). In B the average change in selection rate for each antibiotic compared to LB control is plotted against the average increase in CRISPR-Cas proportion in evolution experiments at 3 d.p.i. (data from Figure 1). with error bars indicating s.e.m.. There was no significant correlation between evolved proportion of CRISPR-Cas clones and CRISPR-Cas selection rate either in the absence of phage (Pearson's product-moment correlation, $t_6$=0.038, $p$=0.97 $R^2$=0.0156) or in the presence of phage ($t_6$=1.05, $p$=0.34, $R^2$=0.39).

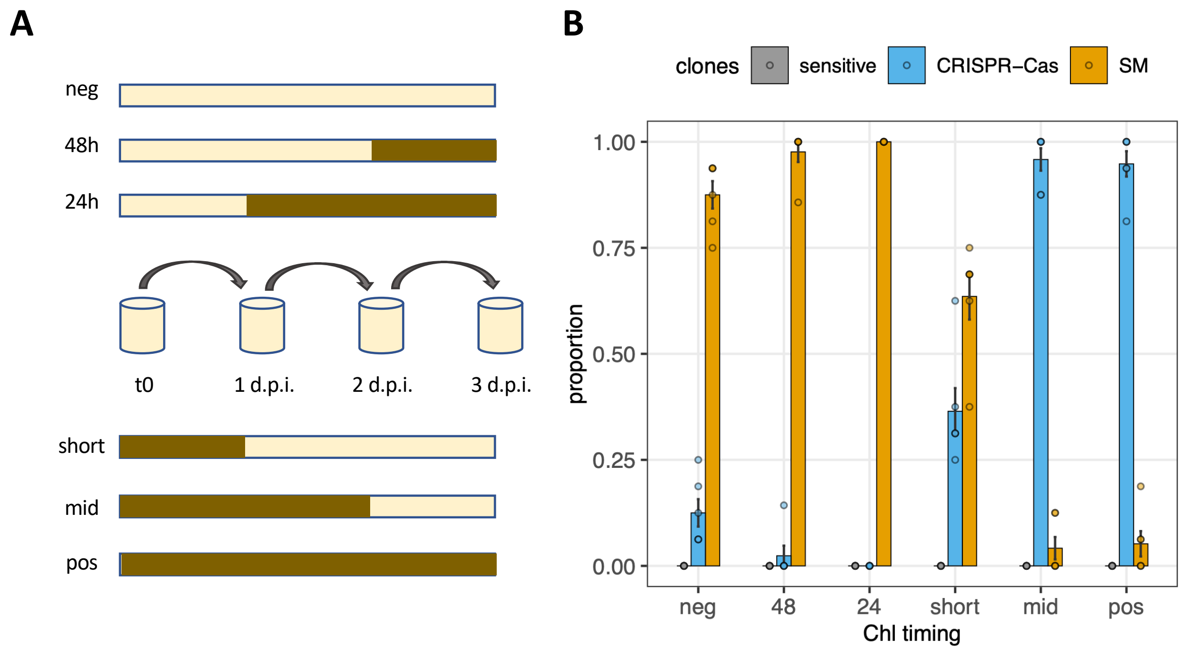


**Figure S9: Effect of the timing of chloramphenicol exposure on CRISPR-Cas evolution.** A shows the experimental treatments varying exposure to 25 mg/L Chl, with dark brown segments representing times in which cultures were exposed to Chl. B shows the proportion of clones with each resistance phenotype at 3 d.p.i.. Bars and error bars show mean ± s.e.m., and individual biological replicates are plotted as dots.

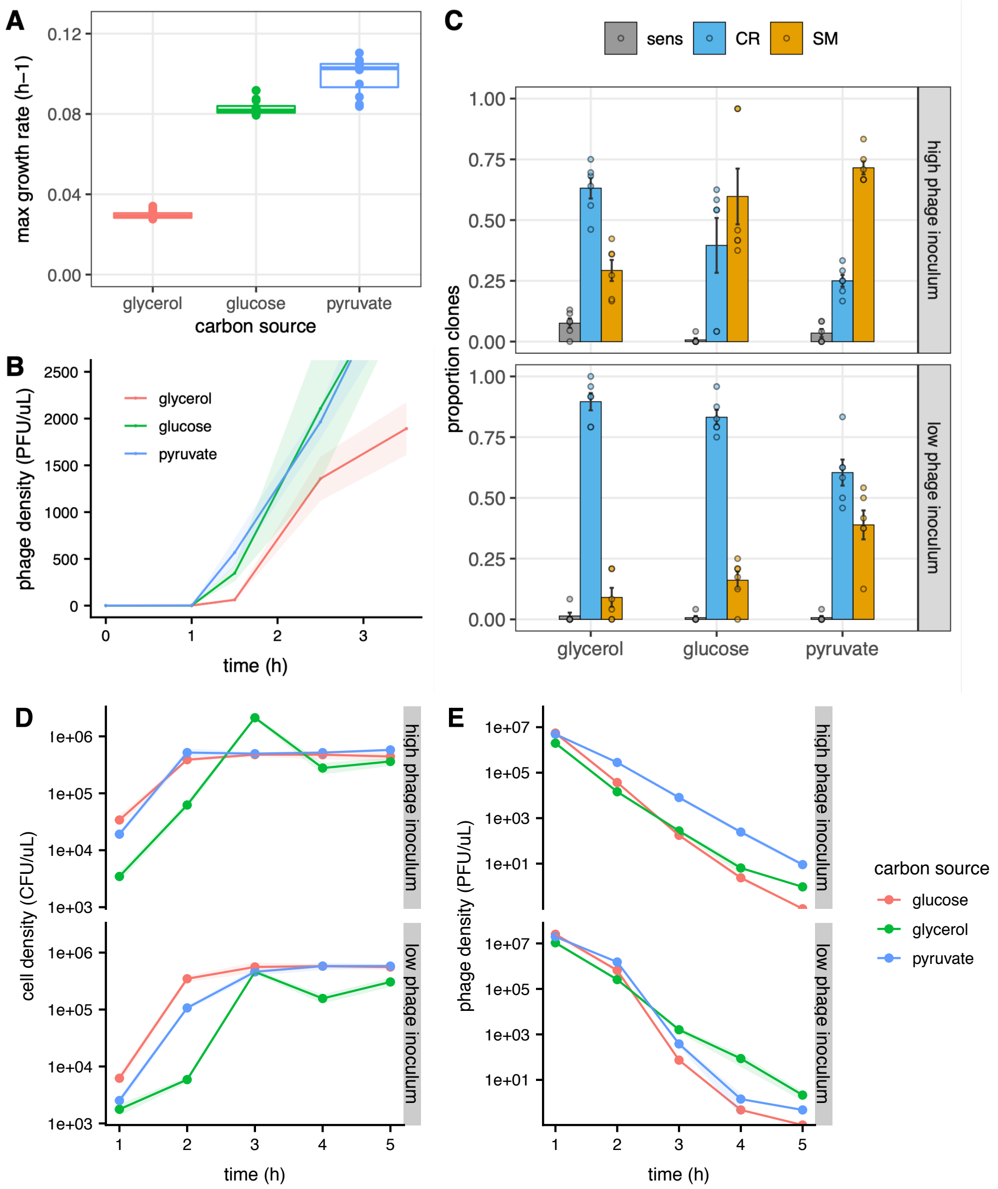


**Figure S10: Carbon sources causing slow growth rate and delayed phage production also promote CRISPR-Cas evolution.** A shows the maximum growth rate from PA14 grown in M9 + 40mM of three carbon sources. The centre value of the boxplots shows the median, boxes the first and third quartile, and individual data points are shown as dots (N=12). In B, phage one-step growth assays are shown. Colour indicates the carbon source present, lines and shaded area are respectively the mean and s.e.m. (N=4). In C, the C shows the proportion of sensitive, CRISPR-Cas and SM clones at 5 d.p.i. is shown. Bars and error bars show mean ± s.e.m., and individual biological replicates are plotted as dots (N=6). In D and E, cell density (D) and phage density (E) are shown over time with color indicating the carbon source in M9 minimal medium. Lines and shaded area show respectively mean and s.e.m. (N=6).
